# Cell extract-based in vitro DNA replication enables sustainable cell-free gene expression

**DOI:** 10.1101/2024.10.01.616023

**Authors:** Xiao Zheng, Wenli Gao, Wan-Qiu Liu, Yufei Zhang, Yicong Lu, Shuhui Huang, Xiangyang Ji, Yifan Liu, Shengjie Ling, Jian Li

## Abstract

Cell extract-based cell-free gene expression (CFE) systems break cellular boundaries and create emerging platforms for synthetic biology applications. However, these systems lack intrinsic DNA replication capability, thereby limiting sustained transcription and translation in a cell-free environment. Here, we introduce a LoopReX system based on *E. coli* CFE by integration of the DNA replication machinery, enabling iterative in vitro DNA replication and sustainable cell-free gene expression. We establish LoopReX by integrating phi29 DNA polymerase into the crude lysate-based *E. coli* CFE system, allowing for coupled DNA replication and protein expression. We elucidate the synergetic mechanisms between T7 RNA polymerase and phi29 DNA polymerase in facilitating in vitro DNA replication. Additionally, we optimize LoopReX reaction conditions by machine learning strategies. Finally, we demonstrate that the enhanced LoopReX system achieves highly efficient DNA replication and protein production in an iterative, scalable, and sustainable manner. We anticipate that LoopReX will significantly advance the application of CFE in synthetic biology.

## Introduction

Life (or a cell) exists primarily to ensure survival rather than to express large quantities of proteins unrelated to that purpose^1^. In living cellular systems, the maintenance of basic metabolism and the expression of heterologous proteins are mutually constrained, often leading to suboptimal efficiency for both processes. To meet the growing demand for protein products, developing more controllable non-living protein expression platforms is essential. In this context, cell-free gene expression (CFE) systems represent the closest approach to a fully controlled in vitro protein production platform. Crude extract-based CFE systems harness the intracellular biocatalysts and machinery (e.g., aminoacyl-tRNA synthetases, RNA polymerases, and ribosomes, etc.) to perform transcription and translation (TX-TL) with the supplementation of necessary substrates, cofactors, and target gene templates^2-5^. While CFE systems exhibit strong protein expression capabilities without maintaining cellular primary metabolism, they lack the ability to replicate themselves, particularly with respect to the supplemented DNA templates^2^. This key limitation renders CFE an unsustainable platform for protein production due to the absence of the essential central dogma system.

Recently, in vitro transcription–translation-coupled DNA replication (TTcDR) system was constructed by introducing DNA replication machinery into the PURE (Protein synthesis Using Recombinant Elements)-based CFE system^6-13^. Specifically, phi29 DNA polymerase (phi29DNAP) is often used in TTcDR for DNA replication due to its high processivity, high fidelity, and initiation without a primase^14^. By introducing Cre recombinase, TTcDR enables the adaptive evolution of artificial genomic DNA, generating performance-enhanced DNA polymerase and recombinase^12^. Moreover, self-replication and expression of large synthetic genomes (>116 kb, close to the predicted genome size required to encode a minimal cell) has also been achieved in a cell-free TTcDR system, demonstrating the potential toward the construction of artificial cells in the future^7^. These above achievements demonstrate the success to establish DNA replication-coupled TX-TL systems using PURE, yet efforts to develop crude lysate-based CFE systems for the coupling of DNA replication remain underdeveloped.

Due to the open nature and easy preparation of CFE systems, this cell-free biotechnology has been widely designed for various applications, ranging from fundamental research to potential industrial biomanufacturing^2,5,15,16^. For example, CFE systems have been utilized for the production of high-value compounds^17-22^, prototyping metabolic pathways^23,24^, screening protein variants^25,26^, detecting small molecules and heavy metals^27,28^, synthesizing clinical therapeutics^29^, designing diagnostics^30^, and constructing synthetic cells^31^, among others. These impressive applications have driven the development of novel and function-enhanced CFE systems^32-37^. However, to our knowledge, no attempts have been made to create TTcDR systems relying on the widely used, crude extract-based CFE platforms.

To address this opportunity, we report a LoopReX (i.e., a loop of DNA replication and protein expression) system for in vitro DNA replication, enabling iterative and sustainable cell-free gene expression (**Fig. 1**). The foundational principle is that we can integrate the phi29DNAP machinery into the cell extract-based CFE system to achieve a minimal in vitro central dogma consisting of DNA replication and TX-TL. We choose phi29DNAP because it possesses exceptionally high processivity (up to 70 kb per single synthesis event) and high DNA synthesis fidelity due to its 3′-5′ exonuclease proofreading activity^38,39^. However, some issues related to the working mechanism of phi29DNAP in cell-free conditions (e.g., the PURE-based TTcDR system) still remain to be elucidated. For example, the initiation process of DNA replication is unclear without the involvement of a terminal protein and supplemented primers^6,8^. Using LoopReX, we aim to investigate the initiation mechanism of phi29DNAP in our cell-free environment. Then, we optimize the LoopReX system by using machine learning to identify the key parameters for the coupling process of DNA replication and gene expression. With the optimized LoopReX system, we achieve higher DNA replication efficiency and faster protein expression compared to the pre-optimized system. Finally, we carry out scalable iterative cell-free reactions, enabling sustainable and high-yield protein production (approximately 3-4 times improvement compared to the control systems). Looking forward, our LoopReX system will provide a robust platform not only for fundamental research such as construction of artificial cells but also for applied biotechnology applications, for instance, biomanufacturing of valuable biomolecules and (bio)chemicals.

**Fig 1.**
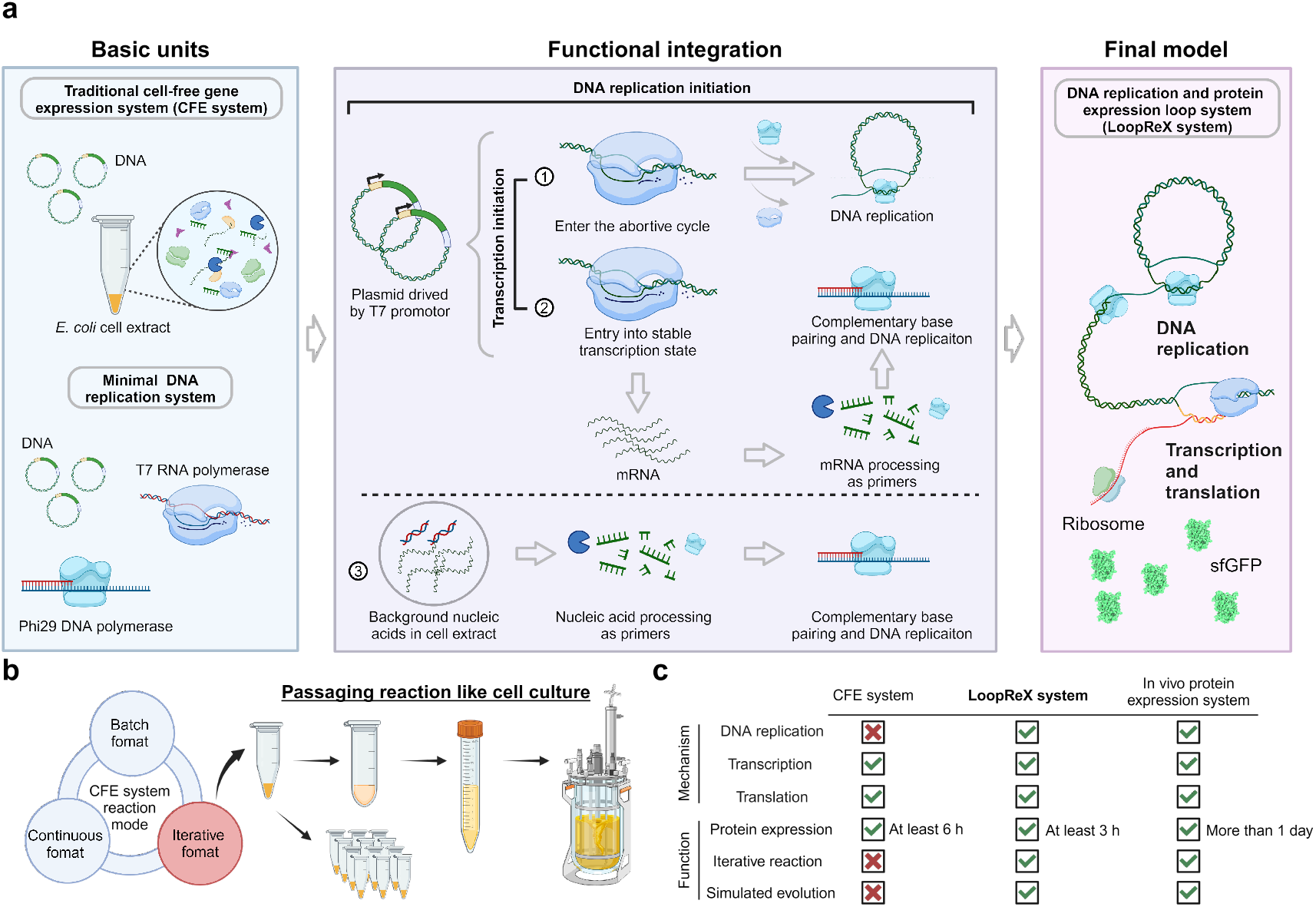
Crude extract-based CFE systems incorporating a DNA replication mechanism for iterative DNA replication and protein expression. **a**. Composition of the LoopReX system and a schematic of DNA replication initiation within the LoopReX system. **b**. Schematic of the iterative reaction mode in the LoopReX system. **c**. Comparison of the mechanism and function between conventional CFE systems, in vivo protein expression systems, and the LoopReX system.

## Results and Discussion

### Establishing LoopReX for the coupling of DNA replication and gene expression

To begin, we mixed M13mp18 single-stranded DNA (ssM13) with a replication reaction buffer, including specific primers, dNTPs, and purified phi29DNAP, as described previously (**Supplementary Fig. 1**)^40^. We incubated this mixture for 1 h at 30°C and verified that an average of 3.6-fold rolling circle amplification (RCA) product was successfully synthesized (see **Methods** for details, **Supplementary Fig. 2**). In the process, through agarose gel electrophoresis of the RCA products, we observed some diffuse bands smaller than the template size, indicating the presence of DNA degradation caused by the inherent DNA degradation activity of phi29DNAP (**Supplementary Fig. 2b**)^41,42^.

To further investigate the RCA process mediated by phi29DNAP, we conducted DNA synthesis time gradient experiments, which are analyzed by agarose gel electrophoresis and real-time quantitative PCR (qPCR). The results indicated that the DNA content reached its maximum within 2 h, with a “primer-to-replication product” conversion of 70-90% across three independent experiments (**Supplementary Fig. 3**). However, due to the exonuclease activity of phi29DNAP, some primers were consumed, suggesting that actual conversions might be higher than the observed data^40^. At this point, the availability of free primers in the system decreased. After 1 h, the total DNA content started to decline, indicating that DNA amplification was primarily driven by the conversion of primers into replication products, as previously reported^40^. With primers exhausted and pyrophosphate accumulated in the reaction, the DNA synthesis rate of phi29DNAP gradually decreased, while exonuclease activity remained unaffected, resulting in a gradual decrease in DNA content after 2 h^38^. Remarkably, despite DNA content gradually decreased, the length of DNA products continued to increase before stabilizing (**Supplementary Fig. 3**). Additionally, degraded DNA bands progressively shifted to smaller fragments and eventually disappeared. We infer that this phenomenon is due to the longer DNA chains may degrade more slowly due to their complex structures, whereas shorter DNA chains degrade more rapidly. As a result, with the reaction progressed, the total DNA content gradually decreased, with the positions of the long DNA fragments remaining largely unchanged and the bands of shorter DNA fragments progressively fading.

Following the initial experiments, we then introduced the DNA replication machinery into the crude extract-based CFE system. To do this, we mixed ssM13 DNA with a CFE reaction that also included specific primers, dNTPs, and purified phi29DNAP. After 1 h, we observed a 4.7-fold increase in the total DNA amount (**Fig. 2a**). In contrast, the total DNA amount decreased to approximately 30% of the initial substrate (ssM13 DNA) in the control reaction without phi29DNAP. This suggests that free nucleases within the CFE reaction rapidly degrade the substrate DNA, implying that the actual DNA replication yield is likely greater than the observed 4.7-fold improvement. Subsequently, we employed double-stranded circular DNA (pJL1-T7-sfGFP) instead of ssM13 DNA as templates, without adding additional primers, to establish the LoopReX reaction. The pJL1-T7-sfGFP plasmid contains a T7 promoter-driven superfolder green fluorescent protein (sfGFP) coding sequence. This design theoretically permits phi29DNAP to autonomously replicate DNA without the requirement of additional replication primers^7^.

**Fig 2.**
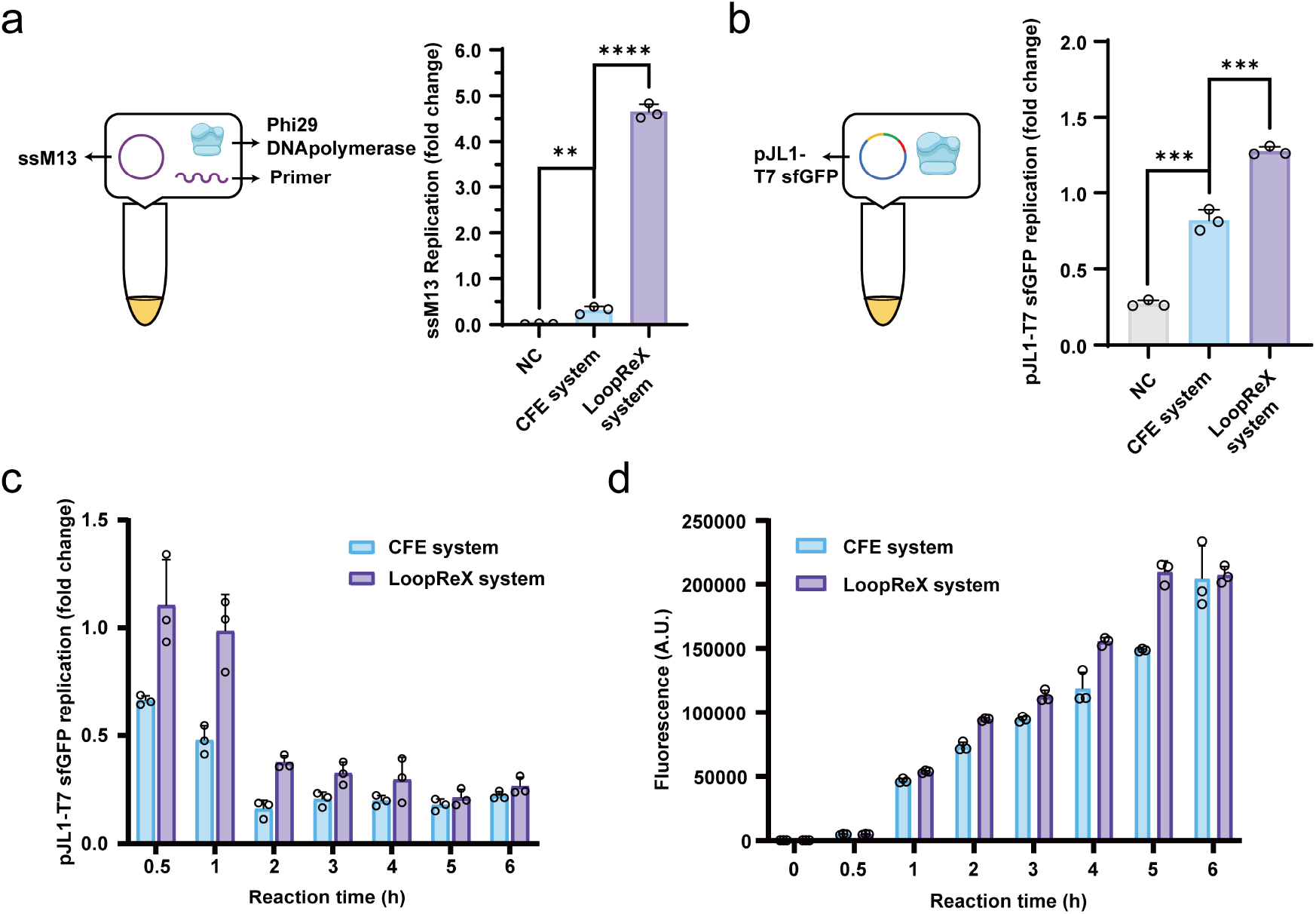
Establishing the LoopReX system for in vitro DNA replication and protein expression. **a**. Replication of ssM13 in the LoopReX with the addition of specific primers after 1 h at 30 °C. Fold changes were determined by qPCR relative to the ssM13 input levels. **b**. Replication of pJL1-T7-sfGFP in the LoopReX without the addition of specific primers after 1 h at 30 °C. **c**. Time gradient reaction of DNA replication in CFE system (blue) and LoopReX system (purple) within 0.5 to 6 h. Note that fold changes after 3 h show no significant difference. **d**. Time gradient reaction of protein expression in CFE system (blue) and LoopReX system (purple) within 0 to 6 h. In panels **(b)** and **(c)**, fold changes were determined by qPCR relative to the pJL1-T7-sfGFP input levels. NC, negative control without gene templates in the CFE system. All data shown representative of three independent experiments (*n* = 3). Statistical significance is indicated as follows: p < 0.05 (*), p < 0.01 (**), p < 0.001 (***), p < 0.0001 (****).

As anticipated, a 1.3-fold increase of the DNA concentration was observed in the replication products compared to the initially added templates (pJL1-T7-sfGFP). However, we observed that the replication efficiency of pJL1-T7-sfGFP was lower than that of ssM13 DNA (**Fig. 2b**). This is expected that double-stranded circular DNA, without added primers, shall undergo helical unwinding, strand separation, and synthesis of replication initiation primers before phi29DNAP can commence replication. In contrast, these prerequisites are present in the ssM13 DNA experiments from the beginning. Furthermore, the double-stranded circular DNA (pJL1-T7-sfGFP) exhibited lower degradation rates compared to ssM13 DNA in control samples without phi29DNAP. This can be attributed to the greater resistance of double-stranded circular DNA to nuclease degradation relative to single-stranded circular DNA.

We then compared DNA contents and protein expression levels in the CFE and LoopReX systems. Our results indicate that the LoopReX system is capable of both DNA replication and rapid protein expression (**Fig. 2c,d**). In contrast to the traditional CFE system, where DNA levels continuously declined, the DNA content in the LoopReX system increased within 1 h and was significantly higher than that in the CFE system within 3 h. After 3 h, DNA levels in both systems showed background values as observed in previous experiments (**Fig. 2b**). Regarding protein expression, the LoopReX system demonstrated accelerated expression rates and reached the plateau phase earlier than the traditional CFE system. This phenomenon is attributed to the increased availability of transcription and translation templates resulting from DNA replication.

Taken together, we demonstrate that: (i) phi29DNAP can degrade DNA within the system, and under identical conditions, high-molecular-weight rolling circle concatemer DNA degrades at a slower rate compared to low-molecular-weight DNA; (ii) phi29DNAP is functional within the conventional CFE system, and free nucleases present in the CFE environment degrade circular DNA; and (iii) the LoopReX system, which introduces phi29DNAP into the CFE system, can replicate DNA and enables faster protein expression than the traditional CFE system. However, the mechanism of spontaneous replication in phi29DNAP assays without the addition of primase and replication initiation primers remains unclear. Thus, elucidating this mechanism is crucial for further optimization of the LoopReX system.

### Elucidating the synergetic mechanism of T7RNAP and phi29DNAP on DNA replication

In 2015, Sakatani et al. reported a TTcDR system based on the PURE system and phi29DNAP^8^, which hypothesized that RNA transcribed by T7 RNA polymerase (T7RNAP) could sever as a primer for DNA replication. To determine if this phenomenon occurs in our LoopReX system, we prepared cell extracts lacking T7RNAP by culturing cells in the absence of the inducer IPTG. The LoopReX system configured with cell extract lacking T7RNAP (named “-T7pol LoopReX”, Bar 3 in **Fig. 3a**) exhibited a significant decrease in DNA replication activity compared to the original LoopReX system (Bar 2 in **Fig. 3a**). Subsequently, equal amounts of purified T7RNAP were added to both the normal LoopReX and -T7pol LoopReX systems. Addition of T7RNAP resulted in an improvement in DNA replication efficiency in both systems, with a particularly enhanced effect in the -T7pol LoopReX system (Bars 3 and 4 in **Fig. 3a**). However, with more T7RNAP added to the -T7pol LoopReX system, the DNA replication efficiency showed a slightly declining trend (Bars 5 and 6 in **Fig. 3a**). In summary, DNA replication efficiency is indeed directly influenced by the concentration of T7RNAP, and T7RNAP and phi29DNAP could form a minimal replication unit in our LoopReX system.

**Fig 3.**
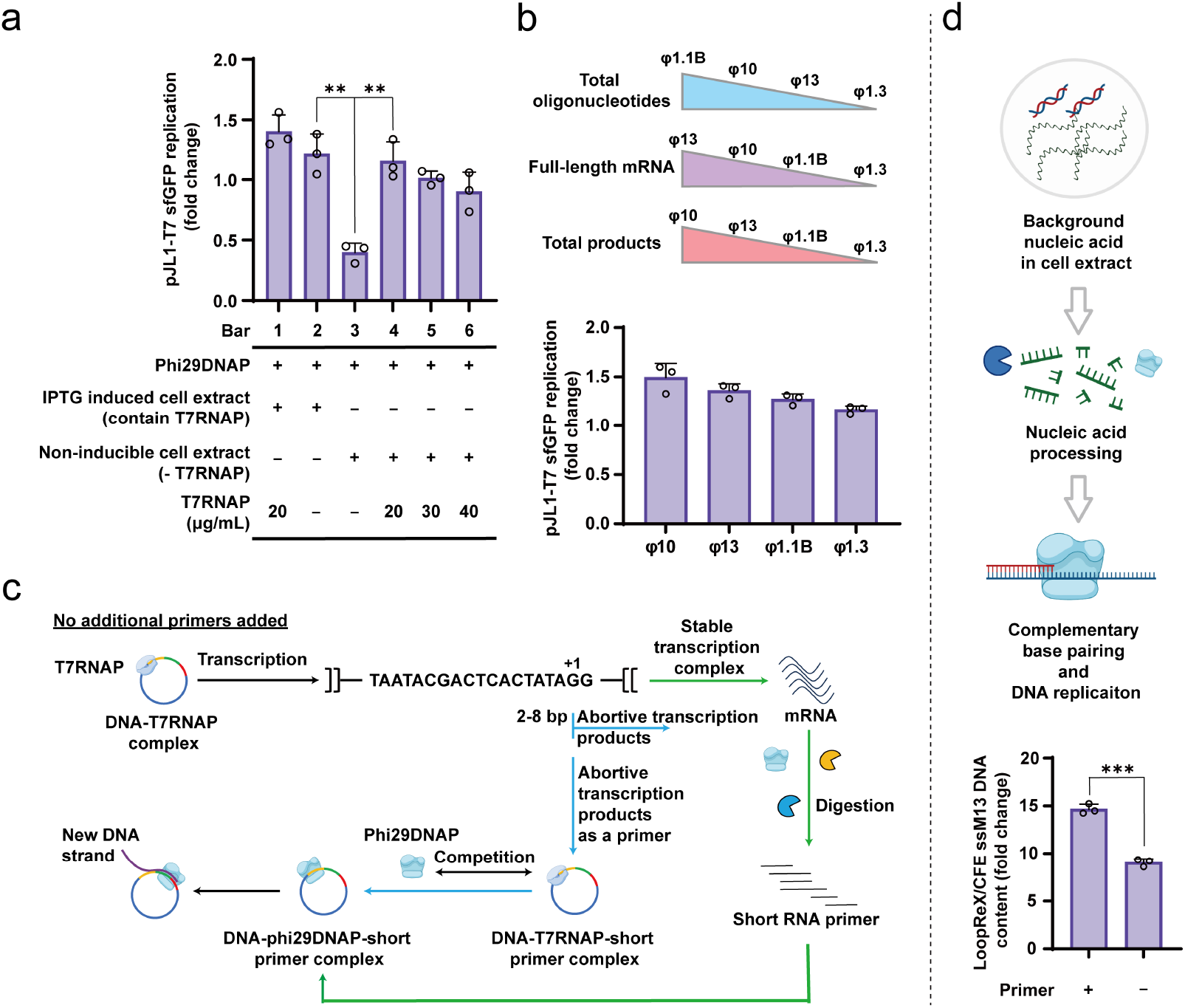
DNA replication initiation in the LoopReX system mediated by T7RNAP at the T7 promoter site. **a**. Replication of pJL1-T7-sfGFP in the LoopReX with different T7RNAP concentrations. **b**. Replication of plasmid driven by different T7 promoters in the LoopReX system. Schematic representation of the preference of the selected promoters for synthesis of oligonucleotides and mRNA, and the total synthesis efficiency. The oligonucleotides represent abortive transcription products of 3-8 nucleotides, with the total product defined as the sum of oligonucleotides and mRNA. **c**. Schematic of replication initiation primers provided by abortive transcription products (blue route) and mRNA (green route). **d**. Replication of ssM13 in LoopReX without the addition of specific primers after 1 h at 30 °C. Fold changes were determined by qPCR relative to the DNA level in the CFE system after 1 h at 30 °C. Schematic representation of the replication initiation primers provided by the free nucleic acid strand within the cell extract. In panels **(a)** and **(b)**, fold changes were determined by qPCR relative to the pJL1-T7-sfGFP input levels. All data are presented as mean±s.d. All data shown representative of three independent experiments (*n* = 3). Statistical significance is indicated as follows: p < 0.05 (*), p < 0.01 (**), p < 0.001 (***).

In the transcription process, T7RNAP often produces two types of products: abortive transcription products (short, non-functional RNA chains) and full-length mRNA (used for protein synthesis). To further validate the origin of DNA replication initiation primers in our LoopReX system, we constructed sfGFP expression plasmids driven by φ13 class III T7 promoter, as well as φ1.1B and φ1.3 class II T7 promoters, replacing the original φ10 promoter used in the pJL1-T7-sfGFP plasmid. Previous studies have indicated significant differences in the relative production of abortive transcription products and full-length mRNA by these promoters compared to the φ10 promoter^43^. To compare their replication efficiency, each of the four plasmids, including the original φ10 promoter plasmid, was employed to construct the LoopReX reaction. The results indicate that, the replication efficiency of the four plasmids followed the order of total transcript product yields (defined as the sum of abortive transcription products and mRNA yields) from their respective promoters (**Fig. 3b**). This suggests that both abortive transcription products and mRNA can serve as primers for DNA replication, and that the phi29DNAP replication machinery exhibits a preference for primers from one of these sources over the other.

Overall, our results reveal that both products of T7RNAP provide sources for the primers required for DNA replication. However, there is an intrinsic distinction between abortive transcription products and mRNA, because mRNA molecues are significantly longer than the abortive products. This length difference suggests that these products likely contribute to DNA replication through different mechanisms, with mRNA requiring further processing to serve as a primer (**Fig. 3c**). Our findings uncover that the T7 DNA replication model provides strong theoretical support for the role of abortive transcription products as primers in DNA replication^44,45^. The genome of bacteriophage T7 exists as a linear double-stranded DNA, with its replication origin located between 14.75% and 15% from the terminus, where situate a pair of tandem promoters, φ1.1A and φ1.1B, devoid of any intervening open reading frame (ORF)^44^. The replication of T7 DNA primarily involves proteins such as T7DNAP, T7RNAP, gene 4 protein (gp4), and DNA binding protein. Interestingly, while the replication fork can still be observed in the absence of gp4 and the DNA binding protein, it fails to appear without T7RNAP, indicating the critical role of T7RNAP in the replication process^44,45^. Sequence analysis of the DNA replication products reveals the presence of a 10-60 bp RNA sequence at the 5’ end, which originates from the abortive transcription products of the φ1.1A and φ1.1B promoters^45^. Notably, previous studies demonstrated that plasmids carrying the φ13 promoter could initiate replication^45^, suggesting that this replication initiation capability is not unique to φ1.1A and φ1.1B, which is further supported by our experimental data (**Fig. 3b**).

In fact, many RNA polymerases are known to produce abortive transcription products, and it has been well documented that numerous DNA replication processes are initiated by RNA primers generated by either RNA polymerase or primase^46^. A hypothesis suggests that the use of abortive transcription products as primers for DNA replication may have played a crucial role in the evolutionary transition from the RNA world to the DNA world^46^. Theoretically, the phenomenon of RNA polymerases producing abortive transcription products likely emerged early in the evolution of the DNA world. In the RNA world, the generation of numerous short, non-functional nucleotide chains by RNA polymerases would have had no biological significance. However, with the advent of the DNA world, these abortive products gained significance by serving as primers for DNA replication. Consequently, DNA replication and RNA transcription underwent co-evolution, leading to the current observation that RNA polymerases play a role in many DNA replication processes. For example, two back-to-back arranged promoters were detected in *E. coli oriC* and mitochondrial RNA polymerase can participate in DNA replication from the light-strand promoter^47-49^. In our model (**Fig. 3c**), when T7RNAP binds to the T7 promoter and produces abortive transcription products, phi29DNAP competes with T7RNAP for the DNA-abortive product (primer) complex^45^. Upon successful recognition and binding to the exposed 3’ hydroxyl group of the abortive product, phi29DNAP facilitates the transition from transcription to DNA replication, thereby initiating rolling circle replication (**Fig. 3c**, blue route). Previously, Okauchi and Ichihashi achieved the co-evolution of the minimal replication unit (consisting of phi29DNAP and T7RNAP) using the TTcDR system through laboratory-simulated evolutionary methods^12^. Their work yielded phi29DNAP mutants with enhanced replication efficiency and a genome with more T7 promoters inserted after evolution. Our data indicate that T7 promoters can serve as replication origins in the genome, which can support their findings that more T7 promoters favor more efficient genome replication.

Unlike abortive transcription products, the involvement of mRNA in DNA replication was an unexpected phenomenon, which has not been reported to our knowledge. We propose that this might be related to the catalytic capabilities of phi29DNAP. Previous studies have demonstrated that phi29DNAP possesses highly active 3’-5’ exonuclease activity, which enables it to cleave both DNA and RNA strands^42^. This cleavage continues until a complementary region with the target DNA, thereby initiating replication^50^. In our proposed model, mRNA may participate in DNA replication through this mechanism. In the LoopReX system, besides phi29DNAP, various endogenous nucleases from *E. coli* cell extracts are present. These nucleases may work in concert with phi29DNAP to cleave mRNA, producing fragments that can serve as primers. These fragments may bind either to newly synthesized single-stranded DNA or within the replication bubble initiated by T7 RNA polymerase, thereby triggering an alternative mode of DNA replication initiation (**Fig. 3c**, green route). Furthermore, the cell extract contains a large amount of nucleotide chains, comprising both DNA and RNA. We propose that these nucleotide chains could also function as replication initiation primers, facilitating the conversion of single-stranded DNA into double-stranded DNA through a mechanism analogous to that using mRNA to provide primers (**Fig. 3d** and **Supplementary Fig. 7**). Our experiments with ssM13 DNA replication, conducted without specific primers, revealed that while the replication efficiency was lower compared to samples with added primers, the nucleic acids present in the cell extract still effectively supported the synthesis of new DNA strands (**Fig. 3d**). This finding suggests that the nascent strands produced by phi29DNAP and T7RNAP can be efficiently converted into double-stranded DNA, which is competent for transcription and translation.

### Machine learning-driven optimization of LoopReX formulation concentrations

In the LoopReX system, T7RNAP is a pivotal component, orchestrating the unwinding of DNA, strand separation, primer generation for DNA replication, and mRNA transcription. Given the central role of T7RNAP, optimizing the whole reaction condition is crucial for enhancing the overall system performance. Several studies have employed single-factor optimization methods to refine TTcDR systems based on the PURE system^8,13^. These optimizations have revealed that certain factors exert differential effects on DNA replication, transcription, and translation^13^. Due to the complicated components in the LoopReX system, single-factor optimization might be inadequate to efficiently optimize the LoopReX system. Therefore, we utilized machine learning to optimize DNA replication and protein expression in LoopReX, conducting an extensive search across millions of conditions. This approach allowed us to identify the optimal parameters for maximizing both processes simultaneously.

To achieve this objective, we constructed XGBoost models to predict protein expression and DNA replication levels, analyzing factors from the CFE standard protocol^51,52^, as well as dNTPs and DTT, to assess their combined influence on both protein expression and DNA replication efficiency (**Fig. 4a**). In the model, 80% of the data was assigned to the training set, while the remaining 20%, unexposed to the training process, was used for testing. Both models achieved an R^2^ of 0.99 in the training set. For the test set, the DNA replication prediction model achieved an R^2^ of ∼0.81, while the protein expression model reached ∼0.88, highlighting the robustness of the machine learning models (**Fig. 4b**). The correlation heatmap also revealed notable differences in factor influence between our LoopReX and the reported TTcDR systems^8,13^, with clear correlations observed between several factors (**Supplementary Figs. 8** and **9**).

**Fig 4.**
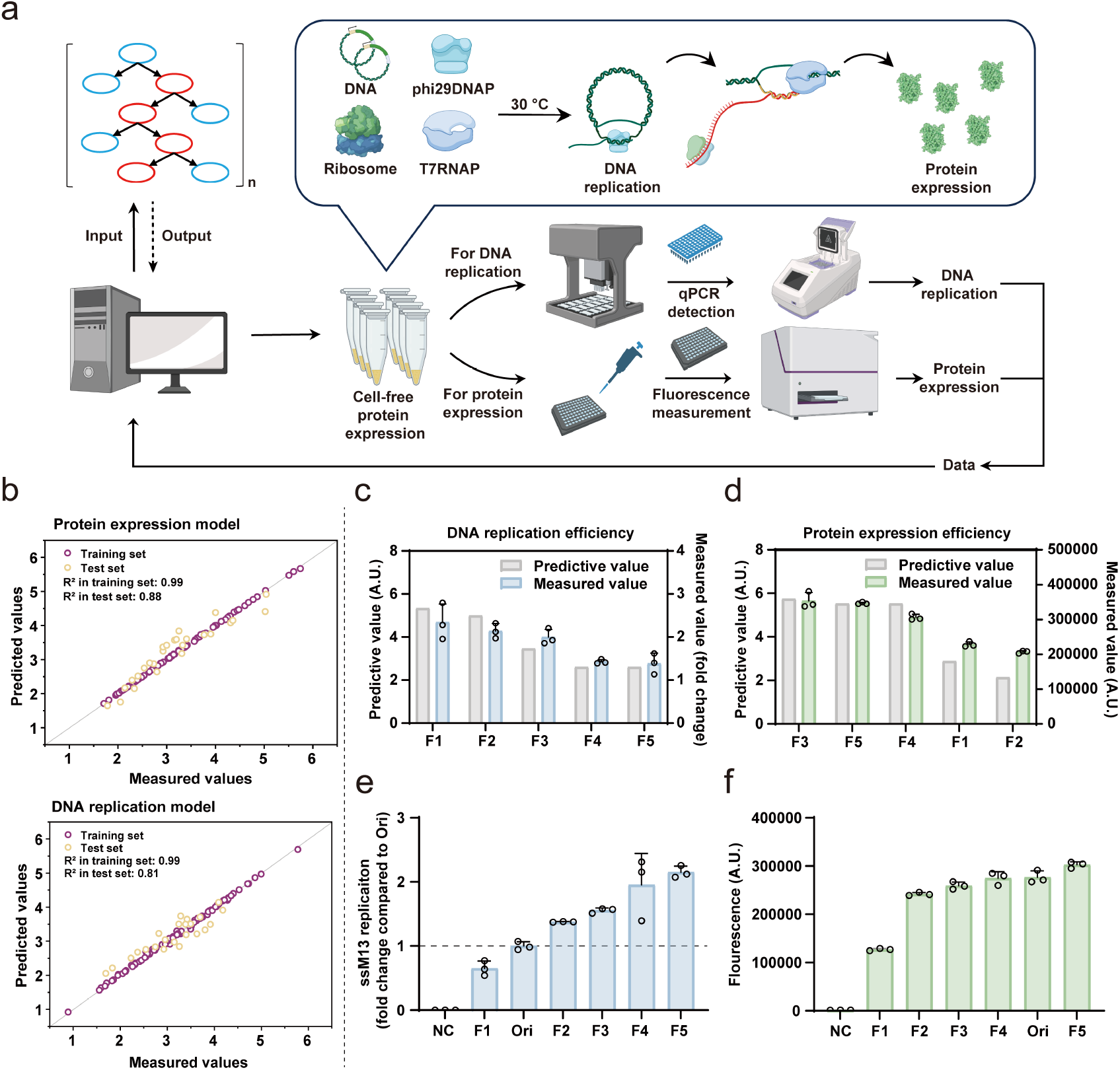
Optimization of LoopReX formulation concentrations via machine learning. **a**. Schematic of the machine learning workflow for optimization of LoopReX formulation concentrations. Experimental procedures for collecting protein expression and DNA replication: DNA content was quantified by qPCR, while protein content (sfGFP fluorescence) was measured using a microplate reader. Since DNA replication and protein expression occur at different time scales, DNA content and sfGFP fluorescence in the LoopReX reaction were collected at 0.5 h and 4 h, respectively. These collected data were used to build models predicting DNA replication and protein expression capacities. **b**. Model performance of protein expression and DNA replication. **c, d**. The pJL1-T7-sfGFP replication (blue) and sfGFP expression (green) under 5 notified compositions and the model prediction results (gray). **e**. The ssM13 replication efficiency of the selected formulation at 0.5 h. Fold changes were determined by qPCR relative to the DNA content in pre-optimized LoopReX (Ori for short) reaction. **f**. The sfGFP expression efficiency of the selected formulation at 6 h, without the addition of phi29DNAP. NC, negative control without gene templates in the CFE system. All data are presented as mean±s.d. All data shown representative of three independent experiments (*n* = 3).

For verification, we selected 5 combinations and conducted LoopReX reaction. For sufficient protein expression, sfGFP fluorescence was collected at 6 h and DNA content were collected at 0.5 h. The results indicated a consistent trend in DNA replication efficiency and protein expression efficiency between the predictions and experiments (**Fig. 4c,d** and **Supplementary Table 4**). To further analyze the above 5 combinations, we investigated DNA replication efficiency using ssM13, which does not contain the T7 promoter; and protein expression efficiency was investigated using pJL1-T7-sfGFP, which was not supplemented with phi29DNAP in the reaction (**Fig. 4e,f**). This approach allows us to separately evaluate the performance of these formulations in DNA replication or transcription-translation. In the absence of protein expression process, the DNA replication efficiency of the formulations from F1 to F5 progressively increases, indicating that in the original LoopReX system, the inhibition of DNA replication by the transcription-translation machinery also gradually intensifies (**Fig. 4e**). On the other hand, the results of protein expression experiments without DNA replication support this observation that the expression efficiency trend across the formulations from F1 to F5 also gradually increases (**Fig. 4f**).

### LoopReX-Opt: Achieving sustainable, iterative DNA replication and protein expression

To enable the optimized LoopReX system to perform sustainable, iterative DNA replication and protein expression, we aimed to identify experimental conditions of both processes. After analysis, we identified the factor concentration combination in the F3 formulation as ideal for meeting the demands of a sustainable iterative system (**Supplementary Fig. 10)**. This balanced F3 formulation (named LoopReX-Opt) supports both DNA replication and protein expression efficiently compared to other formulations (F1, F2, F4, and F5) as well as the pre-optimized system (LoopReX-Ori).

To further investigate the DNA replication and protein expression capabilities of the LoopReX-Opt system, we conducted a series of time gradient experiments. The results indicated that LoopReX-Opt could amplify DNA to approximately twice higher than the initial DNA concentration within 0.5 h, consistent with the efficiency observed during the screening phase (**Fig. 4c** and **Supplementary Fig. 10**), and with no significant degradation observed within 3 h as compared to the control LoopReX-Ori system (**Fig. 5a**). Additionally, we observed a significant improvement in protein expression efficiency using the LoopReX-Opt system. Specifically, the protein expression levels in LoopReX-Opt after 3 h were comparable to those achieved in the LoopReX-Ori system after 8 h and equivalent to the expression levels observed in the traditional CFE system after 14 h, supplemented with the same initial DNA concentration (**Fig. 5b** and **Supplementary Fig. 11**).

**Fig 5.**
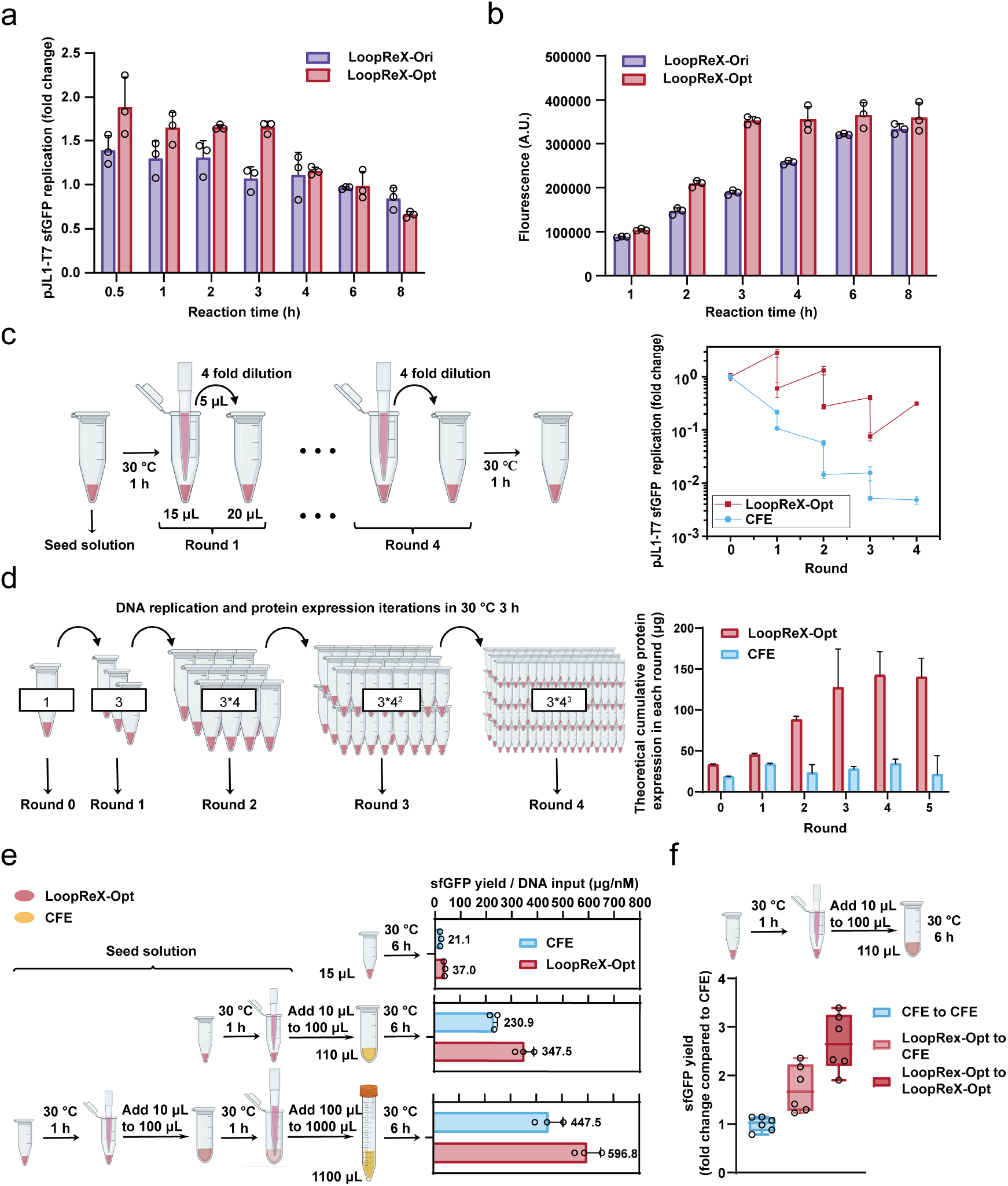
In vitro DNA replication and iterative gene expression based on the LoopReX-Opt system. **a**. Time gradient reaction of DNA replication in LoopReX-Ori (purple) and LoopReX-Opt system (red) from 0.5 to 8 h, with fold changes after 3 h show no significant difference. **b**. Time gradient reaction of protein expression in LoopReX-Ori (purple) and LoopReX-Opt system (red) from 0 to 8 h. **c**. Iterative DNA replication of pJL1-T7-sfGFP through serial transfers in the CFE (blue) and LoopReX-Opt (red) systems. The seed solution, prepared after 1 h at 30 °C, had a standard total reaction volume of 15 μL. For each iteration, 5 μL of this seed solution was transferred into a fresh, plasmid-free reaction system. **d**. Equal-volume saturated iterative gene expression in the LoopReX-Opt system and CFE system. For each iteration, the entire reaction mixture from the previous round was transferred to a fresh, plasmid-free reaction system, following the transfer ratio outlined in panel C (5 μL seed solution into 15 μL total volume). This allowed the first round’s 15 μL to be divided into three tubes, and the second round into four tubes. The bar graph represents the cumulative protein expression within each round of iterative reaction. **e**. LoopReX-Opt based scale-up iterative DNA replication used for protein expression in CFE system. The LoopReX-Opt system was used as a seed solution after 1 h at 30°C, followed by scale-up volume iterations in the CFE system, assessing the impact on protein expression after 6 h at 30°C. Control experiments employed the CFE system as a seed solution without DNA replication capability. The observed increase in protein expression in control experiments is attributed to DNA-to-mRNA amplification, while the difference between experimental and control conditions represents the enhancement in protein expression due to DNA replication. **f**. LoopReX-Opt based scale-up DNA replication and iterative gene expression. Fold changes were determined by sfGFP yields in CFE reactions. In panels **(a-c)**, fold changes were determined by qPCR relative to the pJL1-T7-sfGFP input levels. All data are presented as mean±s.d. All data shown representative of three independent experiments (*n* = 3).

Furthermore, the LoopReX-Opt system had an increased concentration of dNTPs compared to the LoopReX-Ori system (**Supplementary Table 4**). Given the direct impact of dNTPs levels on DNA synthesis efficiency, we sought to raise the dNTPs concentration in the LoopReX-Ori system to match that of the LoopReX-Opt system. As expected, increasing the dNTPs concentration in the LoopReX-Ori system also enhanced DNA replication efficiency, achieving approximately a 2-fold DNA increase compared to the initial DNA concentration (**Supplementary Fig. 13a**). However, when compared to the LoopReX-Opt system, the dNTPs-enhanced LoopReX-Ori system showed slower DNA synthesis. Notably, the LoopReX-Ori system, despite reaching a plateau in protein expression at 3 h, exhibited significantly lower protein expression levels after the dNTPs concentration was increased (**Supplementary Fig. 13b**). These findings suggest that, without adjusting other factors, elevating dNTPs levels can significantly reduce protein expression while also demonstrating that the LoopReX-Opt system has a higher tolerance for dNTPs. This tolerance provides greater flexibility for adding higher dNTPs concentrations for DNA replication. To further evaluate the dNTPs tolerance of the LoopReX-Opt system, we increased the dNTPs levels within the system. We observed that even at 1 mM, the LoopReX-Opt system maintained a 2-fold increase in DNA replication efficiency along with high protein expression levels (**Supplementary Fig. 14a**). However, as the dNTPs concentration continued to rise, both DNA replication and protein expression declined, with protein expression completely lost at 4 mM (**Supplementary Fig. 14b**). Notably, across all concentrations tested, the LoopReX-Opt system consistently outperformed the LoopReX-Ori system in both DNA replication and protein expression. These results further underscore the superior dNTPs tolerance of the LoopReX-Opt system.

As expected, the LoopReX-Opt system proved sustainable for at least four successive iterations of serial dilution, when 5 μL of a 1 h LoopReX reaction was directly transferred into a fresh LoopReX mix (**Fig. 5c**). In contrast, the CFE system did not exhibit this capability. Next, we conducted a series of equal-volume saturated iterative reactions, where the entire reaction mixture from the previous round was used as the seed and combined with an equal volume of fresh reaction mix for the subsequent round, following the aforementioned ratio. We assessed the theoretical protein accumulation by analyzing the minimal unit of each iterative reaction (see **Methods** for details). The results demonstrated that, compared to traditional CFE reactions, the LoopReX system significantly increased protein accumulation with each iteration. By the fifth iteration, 1 nM of initial DNA cumulatively expressed approximately 577 μg of sfGFP in the LoopReX-Opt system, which is 3.6 times higher than that achieved by traditional CFE systems. However, after four iterations, the theoretical protein accumulation plateaued, suggesting that DNA amplification might become less prominent beyond the fourth round (**Fig. 5d**). These findings provide both theoretical and experimental validation for our goal of achieving scalable iterative reactions.

For scale-up iterative reactions, we adapted the overall experimental design to align with cell culture protocols, leading to significant differences compared to equal-volume iterations. First, the aim of the seed solution is to maintain a high DNA level rather than protein levels, so it was transferred after only 1 h of reaction. Second, to ensure complete reaction in the transferred mixture, the reaction time was extended to 6 h. Third, the scale-up cell-free reactions were conducted with shaking to maintain homogeneity. To assess the impact of DNA level differences in the seed solution on subsequent protein expression in scale-up iterative reactions, we conducted both single and double amplification iterative experiments, with all experiments using traditional CFE as a control. In the final round of each reaction, the traditional CFE system was used for protein expression. The results showed that in samples subjected to scale-up iterative reactions, the LoopReX reactions, which incorporate DNA replication, produced more target protein in the final round than the traditional CFE system (**Fig. 5e**). This indicates that the seed solution effectively amplified the DNA during the scale-up process. Notably, the difference between LoopReX and traditional CFE reactions in the double amplification iteration was not significantly greater than in the single amplification iteration, likely due to the suboptimal shaking speed in the 1 mL cell-free reactions (see **Methods** for details). Additionally, in each iteration experiment, the protein expression in the traditional CFE reactions was more dependent on mRNA accumulation.

To demonstrate the advantages of the LoopReX-Opt system, we conducted a single amplification iteration experiment, where the final round of the scale-up iterative process was switched back to the LoopReX-Opt system (**Fig. 5f**). Compared to the control experiments, where both the seed solution and reaction mixture used the LoopReX-Opt system, the samples produced significantly higher protein yields, which is similar to the protein expression observed in the aforementioned 1 mL reaction mixture (**Fig. 5e,f**). This confirms the superior DNA replication and protein expression capabilities of the LoopReX-Opt system.

In summary, to expand the capabilities of CFE systems, enabling them to function as more flexible and versatile platforms for in vitro protein expression, we introduced a DNA replication mechanism into the CFE system. Our goal was to develop the CFE system into a DNA replication-transcription-translation coupled system with the unique capability for iterative reactions. In this study, we introduced the minimal replication unit, composed of phi29DNAP and T7RNAP, into the traditional CFE system (i.e., LoopReX). Time gradient experiments revealed that the LoopReX-Ori system not only maintained strong DNA replication capabilities but also exhibited faster protein expression compared to the traditional CFE system. Subsequently, by altering the promoters on the DNA, we explored the initiation process of DNA replication within the LoopReX system and correlated our findings with evolutionary hypotheses. This work elucidates the mechanism of action of the T7RNAP and phi29DNAP-based minimal replication unit within a CFE system. Considering the complexity of CFE system, we employed machine learning techniques to optimize the LoopReX-Ori system. This approach yielded a range of formulations with F3 showing the optimal balanced efficiency in DNA replication and protein expression.

Finally, we validated the practical applications of the optimized LoopReX system, demonstrating significantly higher protein expression efficiency compared to the Ori system and traditional CFE systems. Remarkably, with only a small amount of DNA added (1 nM final concentration), the LoopReX-Opt system was able to achieve protein expression levels in 3 h equivalent to those of the traditional CFE system after 14 h. Additionally, the LoopReX-Opt system remains robust DNA replication capabilities, making it suitable for DNA replication-related research, including studies on artificial cells, isothermal amplification, and nucleic acid aptamer-based biosensors. Moreover, LoopReX-Opt possesses the ability to perform iterative expression similar to living cells, offering an alternative approach to the traditional CFE reaction modes. Using LoopReX as a seed solution can rapidly amplify DNA, and when the final reaction mixture also utilizes LoopReX, average protein yields can reach up to 5.6 mg/mL within a short period, which is 4.2 times higher than the protein yield reported in standard CFE reaction^53^. This substantial improvement highlights the potential of CFE systems as powerful platforms for both in vitro protein production and broad applications in synthetic biology and biotechnology.

## Methods

### Bacterial strains and media

The *E. coli* DH5α strain was utilized for molecular cloning and plasmid propagation, while *E. coli* BL21 Star (DE3) and its derivatives were employed for in vivo protein expression and preparation of cell extracts. For cultivation, *E. coli* was grown in LB (Luria-Bertani) medium, composed of 10 g/L tryptone, 5 g/L yeast extract, and 10 g/L sodium chloride. For cell extract preparation, cells were cultured in 2xYTPG medium, which includes 10 g/L yeast extract, 16 g/L tryptone, 5 g/L sodium chloride, 7 g/L potassium hydrogen phosphate, 3 g/L potassium dihydrogen phosphate, and 18 g/L glucose, with a final pH of 7.2.

### Construction of expression templates

The gene of phi29 DNA polymerase was codon-optimized and chemically synthesized by GENEWIZ (Suzhou, China). The gene sequences are listed in **Supplementary Table 1**. For in vivo expression, a histidine-tag (6* His-tag) was inserted at *C*-terminus of the gene of phi29 DNA polymerase and cloned into pET28a. The plasmid pJL1-T7-sfGFP (Addgene #69496) was used for DNA replication and protein expression in CFE and LoopReX system. For studying the effect of transcript products on DNA replication, The φ10 promotor of pJL1-T7-sfGFP was replaced by φ13, φ1.1B and φ1.3. The promotor sequences are listed in **Supplementary Table 1**. All primers and plasmids used in this study are listed in **Supplementary Table 2** and **Table 3**, respectively.

### In vivo protein expression and purification of phi29 DNA polymerase

*E. coli* BL21 Star (DE3) harboring the related plasmid was cultivated for protein expression. Seed culture was grown overnight in LB medium. Then, cells were used to inoculate 1 L of fresh LB medium with an initial OD_600_ of 0.05. When OD_600_ reached 0.6–0.8, cells were induced with 0.5 mM IPTG, followed by overnight cultivation at 220 rpm and 16 °C. Then, cells were harvested and the pellets were washed twice with lysis buffer (40 mM Tris, 500 mM NaCl, and 10% glycerol, pH 8.0). After centrifugation at 5000*g* and 4 °C for 10 min, cell pellets were resuspended with lysis buffer and lysed by sonication (50% amplitude, 10 s on/off for a total of 40 min). The supernatant was collected by centrifugation at 20,000*g* and 4 °C for 30 min. The supernatant was collected and repeated once centrifugation. After collection, the supernatant was filtered using a 0.45 μm filter membrane and loaded to a Ni Sepharose 6 Fast Flow column (Cytiva) for purification. Then, the column was washed five times with the lysis buffer containing gradient imidazole concentrations (0, 10, 30 mM, respectively). After the final wash, the target proteins were eluted with elution buffer (40 mM Tris, 500 mM NaCl, 100 mM imidazole, and 10% glycerol, pH 8.0). Afterward, the eluted samples were desalted with the desalting buffer (40 mM Tris, 500 mM NaCl, and 10% glycerol, pH 8.0) using Amicon Ultra-15 centrifugal filters (10 kDa cutoff, Merck) by centrifugation at 5000*g* and 4 °C. The concentrations of purified proteins were determined using a Quick Start Bradford Protein Assay kit (Bio-Rad). The purified products were stored at -80 °C until use.

### Replication assay with single-stranded M13mp18 DNA in reaction buffer

For DNA replication assay based on phi29DNAP, single-stranded M13mp18 DNA (ssM13 DNA, NEB) was hybridized with a 17-mer oligonucleotide primer. The incubation mixture contained, in 20 μL, 50 mM Tris-HCl, 10 mM magnesium chloride, 10 mM ammonium sulfate, 4 mM dithiothreitol, 0.2 mg/mL Recombinant bovine serum albumin, 0.3 mM dNTPs, 5 nM ssM13 DNA, 25 nM oligonucleotide primer (**Supplementary Table 2**) and phi29DNAP. The effect of different phi29DNAP concentrations on DNA replication was examined in this experiment. ssM13 DNA and primers were preincubated at 65 °C for 2 min and subsequently lowered to 25 °C. After incubation for the indicated times at 30 °C, the samples were analyzed by electrophoresis in 1.0% agarose gels alongside DNA length markers (*Trans*2K^®^ Plus II DNA Marker) and real-time quantitative PCR. The results showed that the addition of 10 μM phi29DNAP could produce large-molecular weight rolled-circle concatemer products **(Supplementary Figs. 1** and **2)**. Therefore, subsequent experiments were performed at the concentration of 10 μM phi29DNAP.

### Gel analysis of DNA replication products

Untreated DNA replication samples were directly analysed by neutral agarose gel electrophoresis in 1× TAE (Tris-Acetate-EDTA, gels pre-stained with YeaRed Nucleic Acid Gel Stain), unless otherwise specified. A fraction of the total product remained in the gel pockets, likely due to the large size of some rolling-circle concatemers and/or the potential formation of MgPPi-DNA nanoparticles^54^. In addition, there is another situation that is well judged, protein agglomeration when there is no substrate in the case. To explore the effect of nucleic acids in cell extracts on DNA replication initiation, nucleic acids were purified using a FastPure Gel DNA Extraction Mini Kit (Vazyme) in order to reduce the influence of proteins in the extracts on agarose gel electrophoresis.

### Cell extract preparation

Cell growth, collection, and extracts were prepared as described previously^17,18,22^. In brief, all *E. coli* strains were cultured in 2xYTPG medium. Each 1 L culture was inoculated with an overnight preculture at an initial OD_600_ of 0.05. When the OD_600_ reached 0.6-0.8, 1 mM IPTG was added to induce T7 RNA polymerase expression, and cells were harvested at an OD_600_ of 3.0. The cells were washed three times with cold S30 Buffer (10 mM Tris-acetate, 14 mM magnesium acetate, and 60 mM potassium acetate). After the final wash and centrifugation, the cell pellet was resuspended in S30 Buffer (1 mL/g wet cell mass) and lysed by sonication (10 s on/off, 50% amplitude, input energy ∼600 Joules). The lysate was then centrifuged twice at 12,000*g* for 10 min at 4 °C. The supernatant was flash-frozen in liquid nitrogen and stored at −80 °C until use.

### Cell extract based in vitro DNA replication and gene expression reaction

CFE reactions were performed to DNA replication and gene expression reaction in 1.5 mL microcentrifuge tubes. A standard reaction (15 μL) contained the following components: 12 mM magnesium glutamate, 10 mM ammonium glutamate, 130 mM potassium glutamate, 2.64 mM ATP, 1.89 mM each of GTP, UTP, and CTP, 74.8 μg/mL folinic acid, 375.3 μg/mL of *E. coli* tRNA mixture, 2 mM each of 20 standard amino acids, 0.4 mM nicotinamide adenine dinucleotide (NAD), 0.27 mM coenzyme A (CoA), 1.5 mM spermidine, 1 mM putrescine, 4 mM sodium oxalate, 33 mM phosphoenolpyruvate (PEP), 0.3 mM dNTPs, 1 nM pJL1-T7-sfGFP or 5 nM ssM13 and 25 nM oligonucleotide primer, 27% (v/v) of cell extract. For LoopReX reaction, an additional 10 μM phi29DNAP is required. All reactions were incubated at 30 °C before further analysis of the synthesized DNA and protein. To verify the relationship between DNA replication efficiency and T7RNAP, 27% (v/v) of cell extract without IPTG induction and 20-40 μM T7RNAP are required.

### Relative DNA quantification by qPCR

Fold changes of DNA copy-number relative to input levels (*t* = 0) were measured by qPCR (ChamQ SYBR Color qPCR Master Mix, Vazyme) in a two-steps Real-Time PCR method (CFX96™Real-Time System, Bio-Rad). For each time point, three individual samples were taken and diluted 10,000,000-fold for ssM13 and 1,000,000-fold for pJL1-T7-sfGFP in ddH_2_O, which were further diluted 1:20 in the final qPCR reaction. In order to reduce the adverse effect of the random error of the experimenter’s operation on the machine learning of a large amount of data, the dilution process and qPCR sample loading process were completed by an automated liquid handling workstation (Agilent BRAVO liquid handler). The specific primers for each target amplicon are listed in **Supplementary Table 2**. Melt curve, melt peak and standard curve draw by Bio-Rad CFX Manager 3.1. The fold change is defined as DNA content at that time / DNA input, where “DNA content at that time” and “DNA input” were both detected by qPCR in each independent experiment. All data shown representative of three independent experiments (*n* = 3). The qPCR standard curves involved in this experiment are listed in **Supplementary Figs. 4-6**.

### Protein expression assay

In this experiment, the expression of sfGFP was uniformly used to represent the protein expression level. After CFPS reactions, the fluorescence of sfGFP was measured using a microplate reader (Synergy H1, BioTek). Briefly, 2 μL of the CFPS sample was mixed with 48 μL of nuclease-free water and placed in a 96-well plate with flat bottom. Then, measurements of the sfGFP fluorescence were performed with excitation and emission wavelengths at 485 and 528 nm, respectively. In the final protein expression quantification experiments, the fluorescence of sfGFP was converted to total yield (μg or μg/nM) according to a linear standard curve list in **Supplementary Fig. 12**. This experiment was repeated with three biological replicates. Source data are provided as a Source Data file.

### Factor selection and data processing

The essential functions of cell-free gene expression systems are derived from cell extracts. To sustain high transcription and translation activity outside the cellular environment, traditional cell-free reactions typically incorporate over 50 supplementary components. These components have been shown to influence DNA replication and protein expression through various mechanisms^7,8,13^. Additionally, within the complex milieu of cell extracts, interactions among these factors are likely. Therefore, we collected data on protein expression and DNA replication in our laboratory involving 16 factors in the standard protocol of CFE, including magnesium glutamate, ammonium glutamate, potassium glutamate, ATP, GTP, UTP, CTP, folinic acid, *E. coli* tRNA mixture, nicotinamide adenine dinucleotide (NAD), coenzyme A (CoA), spermidine, putrescine, sodium oxalate, dNTPs, and dithiothreitol (DTT). To reduce noise and errors in the experimental data and improve the accuracy and reliability of the analysis, the Robustness Analysis method (Coefficient of Variation, CV) is used. 126 sets of effective data related to protein expression and 115 sets of effective data related to DNA replication were obtained, respectively.

Furthermore, for parallel experiments in the collected data, Z-score filtering was applied to detect and remove outliers from the experimental data. A threshold of Z=1.5 was set, and data points with Z-scores exceeding 1.5 or less than -1.5 were filtered out. The step of computing the Z-score for each data point are as follows:

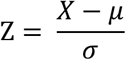

where Z is the Z-score of data X. μ is the mean of data point, and σ is the standard deviation. Then, to make the data more suitable for subsequent analysis, the filtered data was standardized using the Z-Score normalization method, transforming the data to a distribution with a mean of 0 and a standard deviation of 3. To meet specific experimental requirements, the standardized data was further corrected to a range of 0 to 6 to ensure all data points are positive. Since DNA replication and protein expression were theoretically positive results, they were further corrected to a range of 0 to 6 to ensure that all data points were positive values for subsequent model construction.

### Model development

After data processing, we utilized eXtreme Gradient Boosting (XGBoost), an open-source library that efficiently implements the gradient boosting framework, to build the protein expression predictive model and DNA replication predictive model, respectively. Gradient boosting reduces prediction error using a gradient descent algorithm and creates a model composed of multiple weak prediction models, specifically decision trees in this context. During training, new regression trees were added sequentially to minimize residuals (the difference between predicted and actual values), while existing trees remained unchanged to help prevent overfitting. The outputs of new trees were combined with those of existing trees until the loss was minimized to a certain threshold or the specified limit on the number of trees was reached. In this study, XGBoost regressor, implemented through Python code, was trained for each predictor and for the full set of data to learn and understand how many concentrations of factors can influence the efficiency of protein expression and DNA replication. All the data was divided at random into training and test sets with the ratio of 8:2. Only the training set was applied in model training and hyperparameter tuning. To improve the robustness of the model, five-fold cross-validation was used to optimize the model hyperparameter using grid search over the range of possible hyperparameter values. The test set was reserved for testing the performance of the models. Prediction performance was measured using R2, the higher value of R2, the more accuracy of the model.

In this study, we employed machine learning to model DNA replication and protein expression within the LoopReX system, allowing us to identify conditions that optimize both processes simultaneously. Specifically, to find the best conditions for improving protein expression and DNA replication efficiency, we traversed millions of combinations of conditions across the data range. This exhaustive search was designed to rapidly determine the conditions that maximize protein expression and DNA replication efficiency.

### Iterative in vitro DNA replication and gene expression

To validate the DNA iterative replication capacity of the LoopReX-Opt system, we prepared a reaction using the Pr1 formulation according to the standard CFE protocol (15 μL, named standard LoopReX-Opt protocol), incubating it at 30 °C for 1 h as the seed solution. Subsequently, 5 μL of the seed solution was transferred into a fresh standard LoopReX-Opt reaction (15 μL), bringing the total volume to 20 μL, and incubated again at 30°C for 1 h for the first iteration. This process was repeated, and DNA replication capacity was sustained through the fourth iteration.

For equal-volume saturated iterative reactions, the entire reaction mixture from the previous round was used as the seed and combined with an equal volume of fresh reaction mix for the next round, following the previously mentioned ratio. Since LoopReX reaches peak protein expression and maintains high DNA levels within 3 h, the iteration time was set at 3 h. Briefly, after a 15 μL standard LoopReX reaction incubated at 30°C for 3 h, three aliquots of 5 μL were transferred into three fresh 15 μL reactions to begin the first iteration. Starting from this first round, each reaction has a total volume of 20 μL, allowing one 20 μL seed reaction to generate four subsequent reactions. In the second iteration, this process can expand into 12 reactions, and so on. The number of iterations (*n*) and reaction number (*N*) follows the formula:

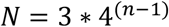

When calculating cumulative protein expression for each iteration, we minimized experimental workload by conducting only the minimal unit for each iteration—transferring one tube of seed solution to 3 reaction tubes in each round. The cumulative protein expression for each iteration follows the formula:

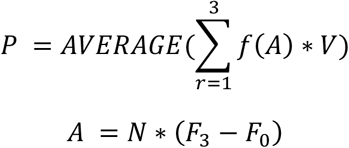

Where:

P is the total cumulative protein expression,

*f*(*x*) is the standard curve of sfGFP,

*V* is the reaction volume,

*r* is the reaction tube in each round,

*F*_3_ is the protein fluorescence value at 3 h and *F*_0_ is the background value,

*N* is the theoretical reaction number.

For scaled-up iterative reactions, we adjusted the overall experimental design to align with cell culture protocols, leading to significant differences compared to equal-volume iterations. Three experimental groups were designed: no iteration, one scaled up iteration, and two scaled up iterations, with approximately 10-fold increases in volume (15 μL, 110 μL, and 1100 μL). Each group’s final round was used for protein expression, incubated at 30 °C for 6 h. The remaining samples served as seed solutions for DNA replication, incubated at 30 °C for 1 h. Fresh reaction volumes were scaled up proportionally to the standard 15 μL protocol. For the first iteration, 10 μL of seed solution was transferred to a fresh 100 μL system and reacted at 600 rpm in a shaker (Thermo-shaker) to enhance oxygenation and maintain homogeneity. For the second iteration, 100 μL of seed solution was transferred to a fresh 1 mL system, reacting at 250 rpm in an orbital incubator (Infors-HT). Due to equipment limitations, the 250-rpm speed may not have been optimal, potentially reducing homogeneity and oxygenation, which could negatively affect the reaction. All control experiments were performed using CFE system as seed or reaction solution. All data shown representative of three independent experiments (*n* = 3).

## Notes

### Competing Interest Statement

The authors have declared no competing interest.

